# Consensus assessment of the contamination level of publicly available cyanobacterial genomes

**DOI:** 10.1101/301788

**Authors:** Luc Cornet, Loïc Meunier, Mick Van Vlierberghe, Raphaël R. Léonard, Benoit Durieu, Yannick Lara, Agnieszka Misztak, Damien Sirjacobs, Emmanuelle J. Javaux, Hervé Philippe, Annick Wilmotte, Denis Baurain

**Author notes:** email list : Luc Cornet Loïc Meunier Mick Van Vlierberghe Raphaël R. Léonard Benoit Durieu Yannick Lara Agnieszka Misztak Damien Sirjacobs Emmanuelle J. Javaux Hervé Philippe Annick Wilmotte Denis Baurain.

## Abstract

**BACKGROUND:** Publicly available genomes are crucial for phylogenetic and metagenomic studies, in which contaminating sequences can be the cause of major problems. This issue is expected to be especially important for Cyanobacteria because axenic strains are notoriously difficult to obtain and keep in culture. Yet, despite their great scientific interest, no data are currently available concerning the quality of publicly available cyanobacterial genomes.

**RESULTS:** As reliably detecting contaminants is a complex task, we designed a pipeline combining six methods in a consensus strategy to assess the contamination level of 440 genome assemblies of Cyanobacteria. Two methods are based on published reference databases of ribosomal genes (SSU rRNA 16S and ribosomal proteins), one is indirectly based on a reference database of marker genes (CheckM), and three are based on complete genome analysis. Among those genome-wide methods, Kraken and DIAMOND blastx share the same reference database that we derived from Ensembl Bacteria, whereas CONCOCT does not require any reference database, instead relying on differences in DNA tetramer frequencies. Given that all the six methods appear to have their own strengths and limitations, we used the consensus of their rankings to infer that >5% of cyanobacterial genome assemblies are highly contaminated by foreign DNA (i.e., contaminants were detected by 5 or 6 methods).

**CONCLUSIONS:** Our results will help researchers to check the quality of publicly available genomic data before use in their own analyses. Moreover, we argue that journals should make mandatory the submission of raw read data along with genome assemblies in order to facilitate the detection of contaminants in sequence databases.

## Introduction

Publicly available genomes are the basic ingredient of numerous studies, from single-gene functional studies to multi-genome phylogenetic inferences. Their quality (e.g., completeness, structural and functional annotation, contamination level) is thus of primary importance. Completeness and annotation have attracted some attention [1–3] but, surprisingly, the issue of contaminating sequences has remained untackled at large scale (i.e., for a whole phylum), despite the well known evidence that contaminants are frequently introduced during experiments [4,5] or stem from natural associations and insufficient purification [6]. Overlooked contaminants may have major detrimental effects on biological conclusions [5]. For instance, the disappearance of two well-accepted monophyletic clades of charophycean green algae (Coleochaetales and Zygnematales) was initially reported [7], but later revealed to be due to cross-contaminating sequences in the dataset made of transcriptomes of green algae and plants [8]. Similarly, another phylogenomic study [9] wrongly concluded to a basal emergence of bilaterian animals, owing to a combination of contamination and taxonomic misidentification [10].

In practice, contaminants arise at different steps, from sampling to sequencing, and anywhere in between [5,11,12]. Yet, for many (microbial) organisms, an aggravating factor is the difficulty to obtain and keep axenic (i.e., pure) cultures [13], explaining why a number of sequenced microbial strains are not devoid of contaminants. While some tools can assess the technical quality of genome assemblies (e.g., QUAST, [2]), or their completeness in terms of gene content (e.g., BUSCO, [1], CheckM [14], ProDeGe [15], acdc [16]) or even their contamination level (e.g., CheckM, [14]), bioinformatic procedures are still needed to recognize and eliminate the contaminating sequences.

As their efficiency ultimately depends on the quality and representativity of the reference databases used for taxonomic classification [3,17], the identification of the genome regions contaminated by foreign sequences (and their elimination) is an important endeavor, both to avoid wrong conclusions and to prevent such genomes from polluting the reference databases, which would lead to misclassifying newly obtained (e.g., metagenomic) sequences.

Recognizing contaminant sequences in genome assemblies, whether prior or after public release, is not a trivial task. When such foreign sequences originate from expected sources (e.g., bacterial cloning vectors, human contamination), they are easy to detect and remove, for example by mapping raw sequencing reads against complete reference genomes of usual contaminant organisms, as part of read quality control before genome assembly (e.g., BBsplit available at https://sourceforge.net/projects/bbmap/). A complication arises with contamination sources closely related to the organism(s) of interest (e.g., human diversity studies or ancient DNA studies). These often require special precautions, both in the laboratory and in downstream analyses [18]. Similarly, parallel processing of multiple organisms, evolutionary related or not, is expected to result in cross-contamination events. Yet, in this case, new sequences can be attributed to their true source based on the comparison of their sequencing coverage across the different samples (Simion et al., 2018, “in press”). With organisms belonging to taxonomic groups for which (high-quality) representative genomes are already available, identifying the genuinely homologous sequences is easy. In contrast, sequences for which there exists no close counterpart in the corresponding reference genomes can be anything from divergent paralogues to new genes, possibly acquired horizontally, or even foreign regions from co-sequenced organisms. Finally, when both the genome under study and its potential contaminants belong to new or scarcely sampled groups, as in the context of large-scale phylogenomic studies, separating the wheat from the chaff can become very challenging [19].

Cyanobacteria, traditionally called blue-green algae, form a large and morphologically diverse group of bacteria [13], which are of primary interest in ecological, palaeobiogeology and evolutionary studies. Hence, the appearance of oxygenic photosynthesis in this phylum had a critical impact on early Earth, its evolution, and on the early biosphere by increasing the level of free oxygen in the ocean and atmosphere, creating new ecological niches [20–22].

Today, they colonize a wide range of illuminated ecosystems, from human-managed to extremophile, and from marine or freshwater habitats (as picoplankton or benthic mats) to hypersaline environments or hot springs, in polar, temperate and tropical regions, with the exception of acidic waters [23]. Usually, contaminants are organisms that live in close proximity to the sequenced organism (i.e., parasites, symbionts, epiphytes) [6,10] or that are simultaneously studied [6,8]. Cyanobacteria are known to excrete different compounds, including polysaccharides, proteins, nucleic acids, osmolytes, in their immediate environment [24,25]. Bacteria from other phyla, such as Bacteroidetes and Proteobacteria, feed on those extracellular productions and thus live in close relationship with Cyanobacteria [24,26,27]. Because of this trophic coupling, the majority of available cyanobacterial strains are not axenic, hence resulting in potential genome contamination. In this respect, we recently noticed that six genome assemblies (GCA_000472885.1, GCA_000817745.1, GCA_000817775.1, GCA_000817785.1, GCA_000817735.1, GCA_000828075.1) presented two or even three copies of many proteins usually encoded by a single gene, one being the likely genuine cyanobacterial protein and the other one(s) being closely related to non-cyanobacterial species (data not shown). These observations prompted us to investigate the issue more thoroughly.

To evaluate the contamination status of public cyanobacterial genome assemblies, we implemented a strategy based on combining evidence across multiple independent methods (**Figure 1**). The rationale was that, given the complexity of the task, no single method would be expected to reach both maximal sensitivity and specificity. Moreover, we were interested not only in quantifying the overall contamination level of cyanobacterial genomes, but also in recognizing and eliminating foreign regions or scaffolds, so as to generate “decontaminated” assemblies. Hence, we first considered a few reliable loci, i.e., genes that have an extremely low probability of horizontal gene transfer (HGT), such as SSU rRNA (16S) and ribosomal proteins. We used RNAmmer/SINA [28,29], an approach based on a reference SSU rRNA (16S) database, and “42”, a BLASTX-based program coupled with a reference ribosomal protein database to detect sequences in cyanobacterial genomes that are phylogenetically related to non-cyanobacterial organisms. Second, we used CheckM [14], a comprehensive package based on the phylogenetic labelling of lineage-specific marker genes, thereby allowing us to probe a larger part of the cyanobacterial assemblies. Third, we explored three genome-wide methods to maximize our detection power. We investigated their parameterization by comparing their results with those based on ribosomal genes, considered as the gold standard. Hence, we tested Kraken [30], a widely used metagenomic classifier based on long (signature) DNA kmers (21–31 nt), against a curated reference database derived from Ensembl Bacteria, and CONCOCT [31], a short DNA kmer (4–6 nt) metagenomic binning package that does not require a reference database, but designed to take advantage of sequencing coverage. As a third genome-wide method, we turned to DIAMOND blastx [32], using the same reference database as with Kraken. Altogether, our analyses allowed us to establish a global ranking of the 440 publicly available cyanobacterial genome assemblies, from the most contaminated to the least contaminated. We conclude that about 20 assemblies are highly contaminated by sequences from foreign phyla, whereas >200 additional assemblies are likely to be at least slightly contaminated. Finally, we provide download links to alternative versions of these assemblies, in which contaminating regions (or whole scaffolds) have been masked.

**Figure 1:**
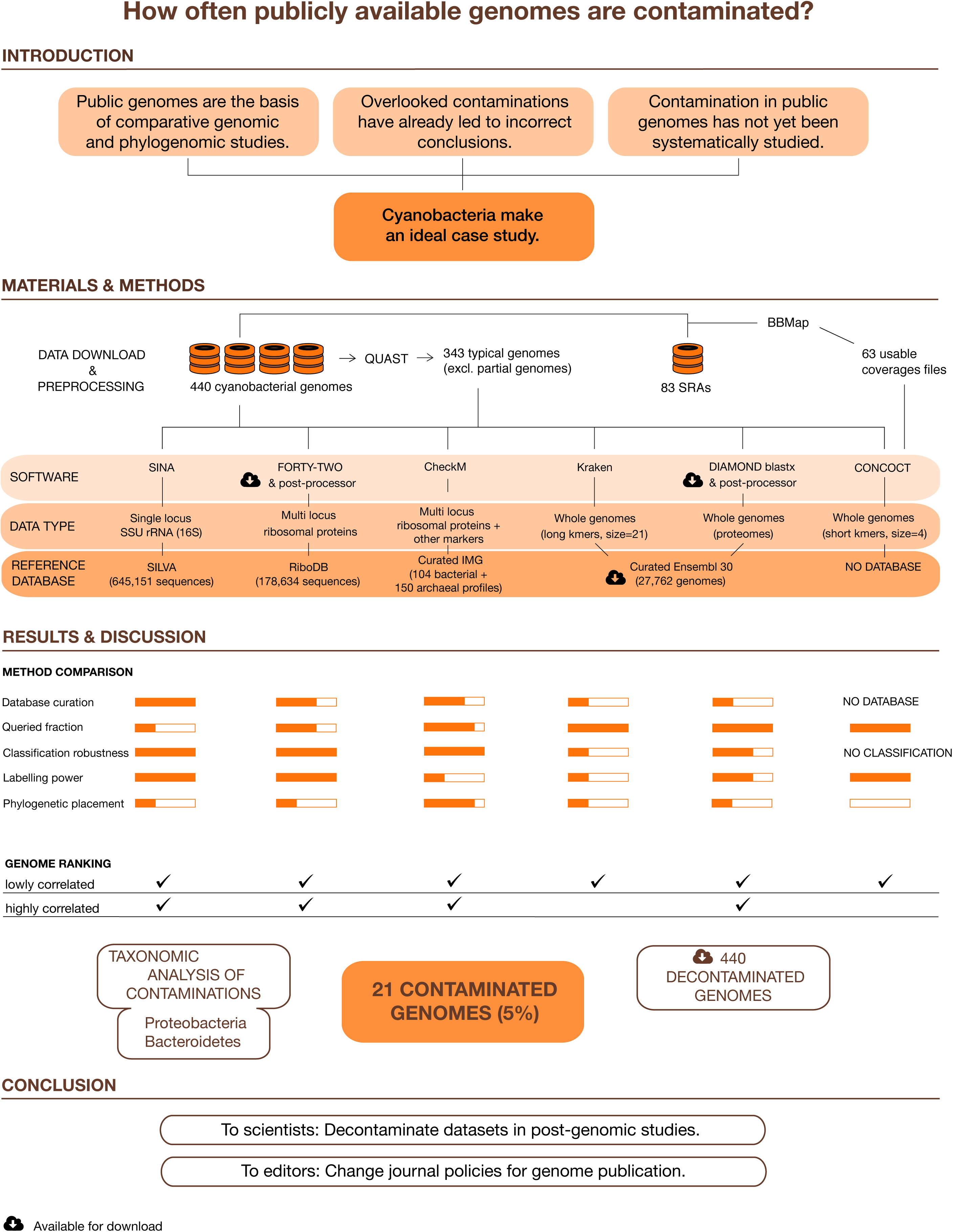
Graphical abstract of the study.

## Results and Discussion

### Properties of the cyanobacterial genome assemblies

We downloaded 440 cyanobacterial genome assemblies, altogether representing the eight orders defined in Komarek et al. [33], as well as the newly erected Gloeomargaritales [34]. These genomes correspond to 421 different organisms, the remaining 19 assemblies being updated versions of existing genomes or independent assemblies (or sequencing plus assembly) of the same strain (**Table S1**). An overview of the morphology, the habitat and the taxonomy of the strains composing our dataset is given in **Figure S1**.

QUAST analyses allowed us to probe the size and the fragmentation level of the assemblies (**Figure 2**). While many unicellular Cyanobacteria have a genome size around 2 Mbp, filamentous, and especially heterocystous, Cyanobacteria are among the bacterial organisms featuring the largest genomes (**Figure 2a**). However, these analyses also revealed that 11 assemblies were <500 kbp, which strongly suggests that they do not correspond to complete genomes, whereas some others are suspiciously large for Cyanobacteria (5 assemblies >15 Mbp, including 2 >68 Mbp but with >90% “N” nucleotides). Regarding the fragmentation level, even if some genomes are assembled into a low number of scaffolds (e.g., one chromosome and a few plasmids), a large part of our dataset consists in genomes represented by more fragmented assemblies, with 57% of the genomes having >20 scaffolds (**Figure 2b** and **Table S1**).

**Figure 2:**
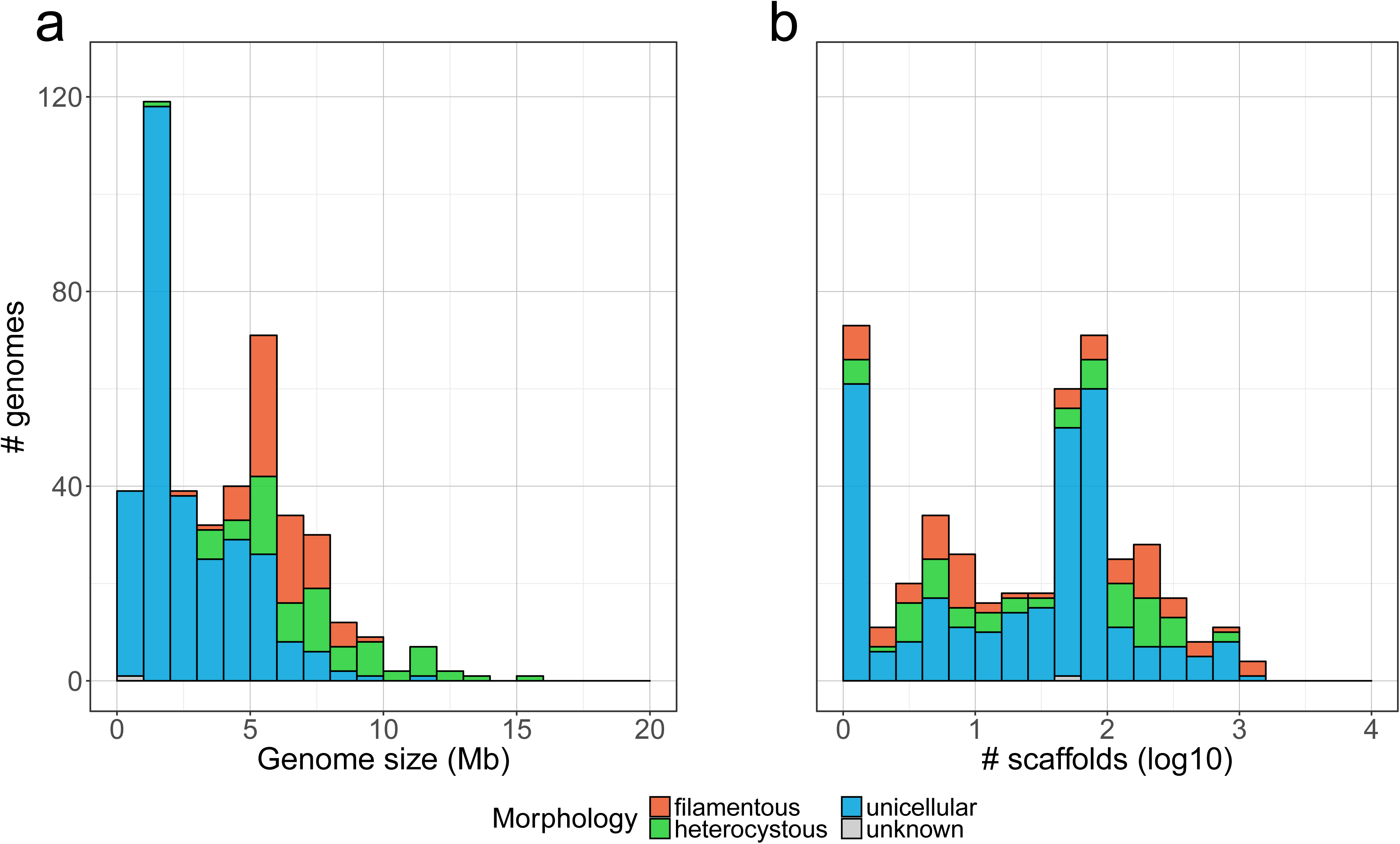
Overview of public cyanobacterial genome assemblies. The 440 strains were classified into four morphologies (unicellular, filamentous, heterocystous, unknown). **a**. Distribution of genome sizes (total length in scaffolds >1000 nt). **b**. Distribution of the numbers of scaffolds (>1000 nt) by assembly. Note the logarithmic scale of the X axis. Details about strain habitats are available in **Figure S1**.

### Ribosomal genes as a first estimator of the contamination level

Since C. Woese and co-workers [35], the SSU rRNA (16S) gene is considered the “gold standard” taxonomic marker. Consequently, large, broadly-sampled and trustworthy reference databases have been available for a long time [36], as well as specialized software to predict (e.g., RNAmmer, [28]) and classify (e.g., SINA, [29]) rRNA genes of unknown origin with both high sensitivity and specificity. Strikingly, 85 assemblies of our dataset appeared devoid of any SSU rRNA (16S) sequence, including 6 of the 13 ultrasmall “genomes” (**Table S1**). Moreover, 4 assemblies contained only unclassified sequences and 1 assembly only one non-cyanobacterial sequence. Among the 350 assemblies featuring at least one cyanobacterial sequence, 15 contained at least one sequence of non-cyanobacterial origin (7 assemblies had 1, 5 had 2, and 3 had 3).

Albeit the presence of one or more foreign SSU rRNA (16S) gene(s) almost certainly reveals a contaminated genome assembly, it only represents a single gene [37], even if it can be found in multiple copies (up to four loci) in Cyanobacteria [38]. To increase the odds of identifying sequences of non-cyanobacterial sources, we turned to a reference database of ribosomal protein genes [39]. While the latter are much less sampled than SSU rRNA (16S) genes, they have the advantage of representing about 50 loci spread over about 10 operons [40]. However, they are not totally immune to HGT (e.g., rps14 [41]). That is why we first inferred phylogenetic trees for all alignments built from the database to identify and remove xenologous reference sequences. Mining of the cyanobacterial genome assemblies against ribosomal protein alignments with our own software “42” (which controls for orthology relationships; available at https://bitbucket.org/dbaurain/42/) showed that 21 assemblies contained at least one ribosomal protein gene of foreign origin (8 in the range [1, 8 proteins], 9 in [16,41] and 4 in [56,80]; **Table S2**). As expected, almost all assemblies featuring at least one foreign SSU rRNA (16S) gene also showed at least one foreign ribosomal protein gene (14 out of 16 assemblies). This allowed us to classify assemblies into three categories, based on the number of ribosomal gene methods identifying contaminating sequences in each assembly: 2 (14 assemblies), 1 (9 assemblies) and 0 (417 assemblies).

### Lineage-specific marker genes as an extended estimator of the contamination level

Even if ribosomal gene methods are sensitive, their power is limited because ribosomal genes represent a small fraction of a cyanobacterial genome (about 0.4–1.0%). Indeed, should ribosomal genes be partially or completely missing from the contaminating fraction of an assembly, it would lead to an underestimation of the contamination level. To mitigate this risk, we turned to CheckM [14], a two-step contamination detection tool based on lineage-specific marker genes, 104 bacterial genes and 150 archeal genes (both including ribosomal proteins, as with “42”).

According to the classification used by Parks et al. [14], CheckM results indicated that 12 cyanobacterial genome assemblies were very highly contaminated (>15%), 2 highly contaminated (>10% to ≤15%), 7 moderately contaminated (>5% to ≤10%), 301 lowly contaminated (≤5%), whereas only 118 assemblies were not contaminated (= 0%). Interestingly, three assemblies (GCA_000472885.1, GCA_000828085.1, GCA_000817785.1) showed a level of contamination >100%, along with a completeness of 100%, thereby suggesting the presence of more than two contaminant organisms in each of them (multiple occurrences of the same markers; **Table 1**). All the 29 assemblies flagged as contaminated by at least one of the two ribosomal gene methods were all also tagged by CheckM. However, 4 assemblies (GCF_000828075.2, GCA_000341585.1, PRJNA165539, GCA_000775285.1) contaminated at >10% in terms of ribosomal proteins were only tagged as lowly contaminated by CheckM (3.41%, 1.96%, 0.88%, 0.84%, respectively). The reason for this discrepancy is likely that CheckM organizes marker genes into sets of collocated genes for estimating contamination. Indeed, genes that reside in close proximity to each other do not provide independent information regarding the overall level of contamination within a genome.

**Table 1:** Global ranking of cyanobacterial genome assemblies. (°) indicates assemblies that are devoid of SSU rRNA (16S) classified as Cyanobacteria; (+) indicates assemblies that are too large (>15,000 kbp); (-) indicates assemblies that are too small (<500 kbp).

| Genome assembly | Assembly properties |  | Ranking results |  |  |  |  |  |  |  |
| --- | --- | --- | --- | --- | --- | --- | --- | --- | --- | --- |
| Accession | Scaffolds.ge.1000.nt | Total.length.ge.1000.nt | rRNA | rprot | CheckM | Kraken | DIAMOND | CONCOCT | rank.avg.6.na | rank.6.na |
| *+GCA_000472885.1 | 974 | 15866152 | 66.67 | 62.5 | 200 | 31.84 | 35.15 | 44.27 | 8.83 | 1 |
| GCF_001482745.1 | 68 | 7647882 | 50 | 40 | 80.53 | 44.93 | 32.99 | 50.41 | 10.25 | 2 |
| GCA_000817745.1 | 296 | 11500044 | 50 | 28.17 | 32.76 | 25.16 | 21.4 | 47.55 | 13.42 | 3 |
| GCA_000817785.1 | 62 | 13096531 | 40 | 41.86 | 104.63 | NA | 20.35 | 38.95 | 16.10 | 4 |
| GCA_000963755.2 | 1320 | 7900996 | 50 | 53.33 | 85.48 | 6.49 | 17.75 | 49.8 | 16.67 | 5 |
| GCF_000963755.1 | 1320 | 7900996 | 50 | 53.33 | 85.48 | 6.49 | 17.76 | 49.13 | 16.83 | 6 |
| GCA_001458455.1 | 336 | 4815140 | 50 | 61.21 | 22.19 | 11.45 | 27.1 | 39.3 | 17.08 | 7 |
| GCA_000817775.1 | 298 | 8799693 | 50 | 13.73 | 23.11 | 21.24 | 15.94 | 36.71 | 18.00 | 8 |
| GCA_000817735.1 | 118 | 11627246 | 33.33 | 44.57 | 44.54 | NA | 18.07 | 34.98 | 19.00 | 9 |
| GCA_000828085.1 | 214 | 10008488 | 50 | 4.35 | 104.63 | NA | 5.51 | 25.69 | 32.90 | 10 |
| °GCA_000634395.1 | 76 | 1282892 | 100 | 35.56 | 11.96 | 11.36 | 41.55 | 21.74 | 33.00 | 11 |
| °GCF_001637395.1 | 420 | 6991351 | 0 | 11.94 | 54.29 | 34.21 | 15.13 | 49.63 | 49.92 | 12 |
| GCA_000828075.1 | 135 | 10627177 | 75 | 44.44 | 5.39 | NA | 12.95 | 14.19 | 51.00 | 13 |
| °GCF_001637315.1 | 171 | 4600567 | 0 | 27.87 | 5.42 | 11.28 | 6.66 | 85.01 | 55.58 | 14 |
| GCF_000828075.2 | 61 | 9468441 | 75 | 40.79 | 3.41 | 0 | 6.23 | 92.28 | 57.00 | 15 |
| GCA_000341585.2 | 10 | 5525469 | 75 | 4.35 | 4.39 | 6.45 | 3.75 | 10.91 | 58.58 | 16 |
| °GCA_000934435.1 | 174 | 8560182 | 0 | 38.46 | 5.53 | 12.63 | 11.15 | 16.43 | 81.42 | 17 |
| GCF_000346485.2 | 27 | 12284271 | 20 | 0 | 2.77 | 0.69 | 1.35 | 86.56 | 93.17 | 18 |
| °-PRJNA165539 | 59 | 327995 | 0 | 10 | 0.88 | 4.45 | 18.13 | 24.59 | 93.83 | 19 |
| GCF_001904775.1 | 44 | 5821893 | 0 | 0 | 1.45 | 13.12 | 2.38 | 96.83 | 98.67 | 20 |
| GCA_000341585.1 | 1354 | 2669044 | 0 | 11.11 | 1.96 | 6.02 | 3.32 | 8.65 | 102.08 | 21 |

### Genome-wide estimation of the contamination level using long DNA kmers

To confirm marker-based results and improve our detection power, we looked for a genome-wide method and used a metagenomic software package relying on the classification of long (21–31 nt) signature DNA kmers, Kraken [30]. Kraken splits genomes into kmers that it organizes in a taxonomic tree that is then queried to classify raw sequencing reads to taxa of increasing ranks, depending on the conservation of their component kmers (see [30] for details). In this work, we “simulated” raw sequencing reads by splitting the cyanobacterial scaffolds into pseudo-reads of 250 nt and fed them to Kraken in order to analyze the taxonomic composition of each assembly. To minimize potential issues due to incomplete or aberrant genomes, we limited the genome-wide analyses reported in this section to the 343 assemblies that were not too small (≥500 kbp) nor too large (≤15,000 kbp) and that contained at least one cyanobacterial SSU rRNA (16S) gene(s). However, all 440 assemblies were eventually analyzed and the corresponding results are shown in **Table S2**, in which those 97 atypical assemblies are denoted by various symbols.

Three variables affect Kraken ability to classify sequences: the kmer size, the reference genome database, and the confidence parameter. To maximize the fraction of classified sequences (including those from evolutionarily isolated assemblies containing many unique kmers), we used a kmer size of 21 nt and built a curated database (27,762 genomes) from the release 30 of Ensembl Bacteria [42]. These important methodological choices are discussed in **Supplemental Note 1** (see also **Figures S2** and **S3a,b**).

The confidence threshold is a parameter meant to adjust the trade-off between specificity and precision (in terms of taxonomic ranks). Because it has a large impact on Kraken behavior, fine-tuning this parameter is a crucial step. To this end, we compared Kraken results obtained on the 343 typical assemblies at three different confidence thresholds (0.02, 0.04 and 0.06) to the results obtained on the same assemblies with the ribosomal gene methods (categories 0, 1 and 2), here considered as the gold standard (**Figure 3a**). We derived a contamination level from Kraken results for each assembly by summing the classifications that corresponded to non-cyanobacterial sources. For some assemblies, a non-negligible pool of classified sequences were actually classified to the high-ranking “Bacteria” taxon (or to the lower “Terrabacteria” taxon). While these sequences were genuinely part of the total (100%), we did not include them in our contaminating fraction, nor in our cyanobacterial fraction and labelled them as “unknown”. For our purposes, lowering Kraken precision thus affects its sensitivity, since a reduced precision increases the share of these classified-yet-unknown sequences.

**Figure 3:**
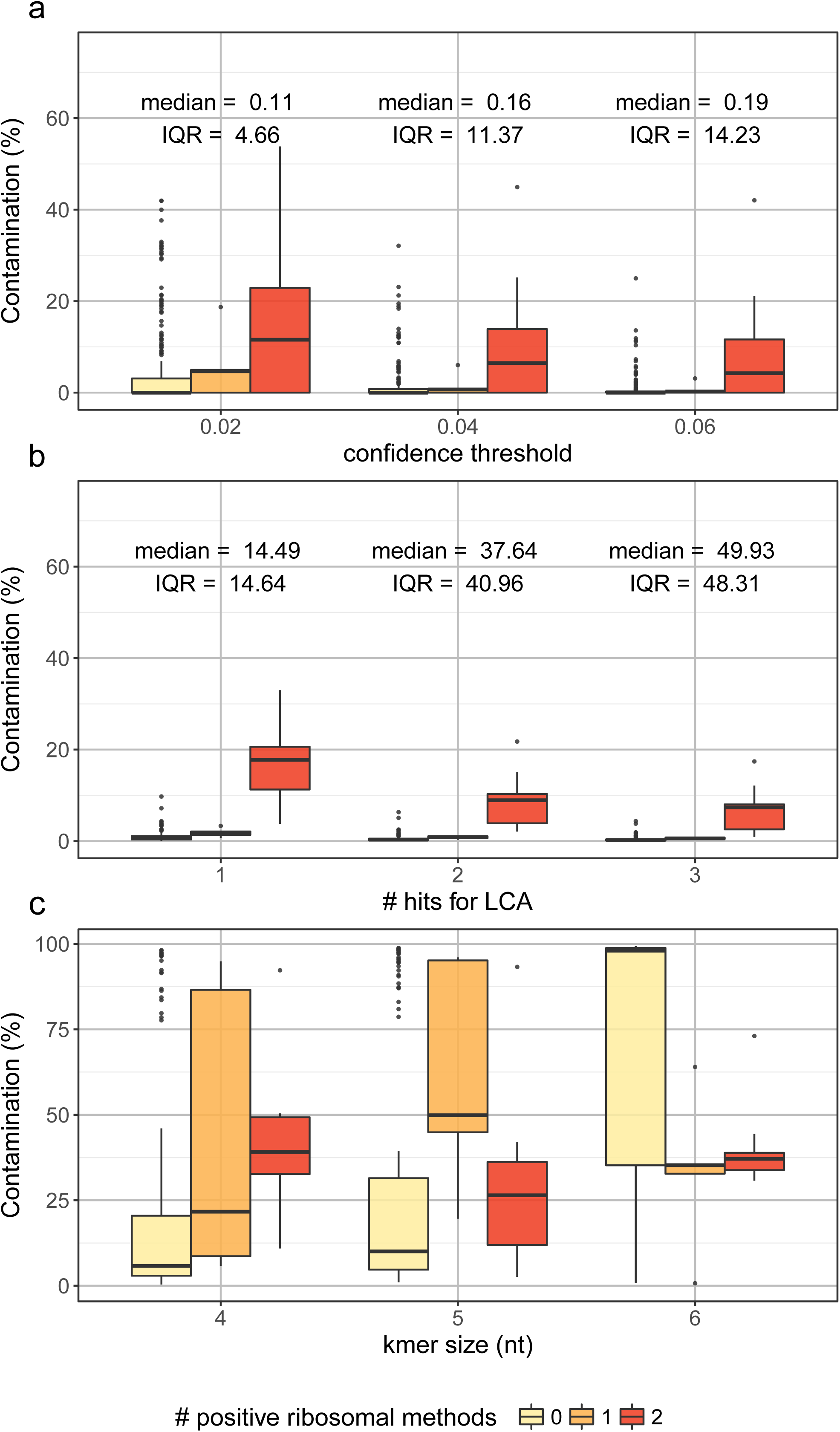
Comparison of genome-wide estimates of the contamination level with ribosomal gene results, considered as the gold standard. Contaminated fractions of the 343 cyanobacterial genome assemblies (expressed in %) were estimated with Kraken using three different confidence thresholds (**a**), or with DIAMOND blastx using three different hit number thresholds (**b**), or with CONCOCT using three different kmer sizes (**c**), then partitioned into three sets of assemblies, based on the number of ribosomal gene methods (SSU rRNA (16S) and ribosomal proteins) identifying contaminating sequences in each assembly (0, 1 or 2). Upper and lower whiskers extend from the hinge to the largest and lowest value no further than 1.5 ^∗^ IQR from the hinge, respectively. Data points beyond the end of the whiskers are outliers and are plotted individually. The values given on top of each series in panels **a** and **b** are the median and IQR for the unclassified fractions across the 343 cyanobacterial assemblies, independently of the ribosomal category.

Based on **Figure 3a**, and with the aim of minimizing the fraction of unclassified/unknown sequences (sensitivity) without wrongly tagging as contaminated too many apparently clean genomes (specificity), we chose to work with the 0.04 confidence threshold to analyze our dataset (**Table S2**). Overall, Kraken analyses provided an independent confirmation that 137 cyanobacterial genome assemblies are contaminated at a level ≥1% (255 at a level >0%). However, a number of these assemblies appear to have contaminating fractions well above the median, despite the SSU rRNA (16S) and ribosomal protein analyses suggesting these genomes are free of contamination (outliers in **Figure 3a**).

### Genome-wide estimation of the contamination level using whole proteomes

Our thorough optimization of Kraken analyses led us to conclude that it is of limited use for our purpose, because it looks for known signature kmers. Given the fast evolutionary rate of nucleotide sequences, genomes of organisms not closely related to any reference genome cannot be classified. Moreover, combining all signature kmers into a single data structure makes it impossible to exclude self-matches when probing an assembly for contamination. This makes Kraken results uninformative for any genome used to build the reference database. To address this problem, we developed a detection protocol based on DIAMOND blastx [32], an ultrafast clone of the BLAST algorithm. In contrast to Kraken, DIAMOND blastx can classify organisms that are distant from all those of the reference database (protein sequences being much more conserved than nucleotide sequences), while still running fast (20,000 times faster than the NCBI BLAST+ implementation) [43]. Besides, it allows skipping self-matches when parsing its output for last-common ancestor (LCA) classification of pseudo-reads. We used the same curated database as for Kraken, except that reference sequences were conceptual translations of the genes instead of whole genomic sequences. Our LCA inference algorithm was inspired from MEGAN [44]. Using a tree-like collapsing of best hits taxonomy down to a bit score threshold expressed as a percentage of the highest bit score, it often allows labelling pseudo-reads based on more than a single best hit.

The better sensitivity of protein searches improved the classification of distant assemblies (**Figure S3c**; slope = -243, Pearson r = -0.76, P-value = 7.92e-35). However, the unclassified fraction of the assemblies was larger with DIAMOND blastx (median = 14.5%, IQR = 14.6%) than with Kraken (median = 0.2%, IQR = 11.4%). This reduced power of DIAMOND blastx is due to the fact that protein-based searches only target coding sequences. Accordingly, the unclassified fraction fits well with the fraction of non-coding sequences in the assemblies (median = 15.9%, IQR = 7.3%).

The potential of DIAMOND blastx was then evaluated exactly as for Kraken, using the 343 cyanobacterial genome assemblies and the ribosomal gene results as the gold standard. A striking difference was that the contamination levels estimated with DIAMOND blastx were much narrower, while still showing a good agreement with ribosomal gene categories (**Figure 3b**). Since the bit-score threshold did not make any obvious difference in our setup, we settled on a value of 5% of the best hit bit score, as in MEGAN (**Table S2**). Even though DIAMOND blastx contamination levels were in line with Kraken levels across all ribosomal gene categories (compare medians of the middle series between panels **a** and **b** of **Figure 3**), all cyanobacterial genome assemblies appeared to contain a non-null fraction of foreign sequences with DIAMOND blastx. In contrast, 181 assemblies were devoid of any foreign sequence with Kraken, including the 170 untestable assemblies (NA in **Table S2**) that are part of its reference database.

As a contribution towards better publicly available genome assemblies, we provide download links to our curated reference database derived from Ensembl Bacteria 30, a post-processor script for DIAMOND blastx output (https://figshare.com/s/344be979f5f2237ad735), and “decontaminated” versions of the 440 cyanobacterial genome assemblies (based on DIAMOND blastx results) (https://figshare.com/s/08193cb71f0a65d99c69).

### Genome-wide estimation of the contamination level using short DNA kmers

As explained in **Supplemental Note 1**, an issue with database-based methods, such as Kraken and DIAMOND blastx, is that assemblies from Cyanobacteria that are isolated from a phylogenetic point of view can be very difficult to investigate due to the proper lack of related reference genomes in the database. To estimate the contamination level while accounting for sequences originating from not yet sampled organisms, we resorted to a program that does not require a reference database: CONCOCT [31]. This software clusters assembly sequences into non-hierarchical groups based on a Principal Component Analysis (PCA) of short (4–6 nt) DNA kmer frequencies. We hypothesized that a high number of CONCOCT groups would hint to a mixture of several genomes into a single assembly. However, we failed to demonstrate any correlation between the number of CONCOCT groups based on kmer frequencies and the DIAMOND blastx contamination level (e.g., for kmer size = 4: Pearson r = -0.04, P-value = 0.31).

By looking at detailed CONCOCT results, we noticed that many cyanobacterial genome assemblies had an overwhelming share of their genomic sequence included in the most abundant CONCOCT group (median = 94.45%, IQR = 15.28%). We thus confidently assumed that this group represented the genuine organism. For practical purposes, we further considered all the other groups as contaminants, which allowed us to compute contamination level estimates by summing the latter fractions. Of course, this is an approximation, because some minor groups might actually correspond to atypical regions of the genuine organism (e.g., repeated elements, plasmids). Nevertheless, when applying this simple metric to tetramer frequencies of the 343 typical assemblies, it performed well in comparison to ribosomal gene methods (**Figure 3c**), in spite of a non-negligible number of assemblies wrongly flagged as highly contaminated (outliers in **Figure 3c**). In contrast, it did not work with pentamers and hexamers, because the most abundant CONCOCT group at these kmer sizes often corresponds to a (much) lower share of the genomic sequences (median = 90.94%, IQR = 21.56% for kmer size = 5 and median = 1.89%, IQR = 62.76% for kmer size = 6).

### Global ranking of cyanobacterial genome assemblies

As the three genome-wide methods had largely confirmed the results obtained with ribosomal genes and lineage-specific marker genes on a subset of 343 cyanobacterial genome assemblies, we decided to use the contamination levels estimated through all six methods to build a consensus global ranking of the 440 assemblies composing our dataset (**Table S2**). Congruence between individual methods and with this global ranking was investigated using Spearman rank correlations (**Table S3**). Ribosomal gene methods correlated weakly when considering either 440 (rho = 0.32/0.37) or 343 assemblies (rho = 0.34/0.35), but worked much better on the top-50 (rho = 0.74/0.79). This is hardly surprising given that, being less sensitive than genome-wide methods for their limited number of interrogated loci, they can only find contaminants in highly contaminated assemblies. On the opposite, the database-free CONCOCT correlated well on the 440 assemblies (rho = 0.63) and this also result held for the 343 assemblies (rho = 0.61). Yet, it did not when focusing on the top-50 (rho = 0.14), as expected from its reasonable but rough metric. The situation for Kraken was similar to that of CONCOCT, with rho = 0.59 (440), rho = 0.59 (343), rho = 0.32 (50), suggesting that in spite of the shortcomings of its monolithic reference database (see **Supplemental Note 1**), Kraken remains useful for real applications. CheckM was the method that correlated the best on average — rho = 0.75 (440), rho = 0.76 (343), rho = 0.73 (50) — followed by DIAMOND blastx — rho = 0.61 (440), rho = 0.68 (343), rho = 0.75 (50). Interestingly, even the best methods had their outliers (e.g., CheckM: GCA_001456025.1 at rank 83, Kraken: GCA_001039265.1 at rank 36, DIAMOND blastx: GCA_000484535.1 at rank 119). This seems unavoidable as pipelines involving several programs and databases are difficult to develop and test exhaustively [45]he contamination level is more valuable than relying on a single approach.

Based on this global ranking, we defined three sets of assemblies. The top-21 (19 different genomes) contains assemblies tagged as contaminated by at least five of the six detection methods (except GCF_001904775.1, which is tagged by four methods) and thus corresponds to those with the highest level of contaminants. Three assemblies (GCA_000472885.1, GCA_000817785.1, GCA_000828085.1) even show CheckM contamination levels >100%, suggesting contamination by more than one organism. For these 21 assemblies, we explored the distribution of contaminants over the genomic scaffolds using DIAMOND blastx (**Figure S4**). In many cases, it is very clear that at least two types of scaffolds co-exist, one corresponding to cyanobacterial segments (in green) and the other to contaminant segments (in red). Such a partition is often reflected in the GC-content of the scaffolds (e.g., GCA_000472885.1, GCA_000817745.1, GCF_001637395.1), and ribosomal gene (both SSU rRNA (16S) and ribosomal proteins) hits nearly always co-localize with the expected type of scaffold (e.g., GCA_000341585.2). This indicates that the different methods are congruent even within a single genome assembly and that their consensus faithfully reflects the underlying contamination level. After the top-21, a second part of the ranking (down to rank 230) contains assemblies that are only slightly contaminated. This is where discrepancies between individual methods are the most common (e.g., GCF_001456025.1 at rank 83, GCA_000291825.1 at rank 227). Finally, the last part of the ranking (210 assemblies) contains genomes that have a very low level of contaminants (generally <1% for both CheckM and DIAMOND blastx estimates).

### Taxonomic analysis of contaminants

According to DIAMOND blastx, the two most frequent contaminant phyla in our top-21 ranking of contaminated assemblies are Proteobacteria and Bacteroidetes (**Figure 4**; see **Figure S5** for a similar analysis for all 440 assemblies). Proteobacteria contaminate 19 assemblies of the top-21 at a level ≥1%, Bacteroidetes contaminate 5 assemblies, whereas Firmicutes and Chloroflexi contaminate 1 assembly. Other contaminants (Acidobacteria, Actinobacteria) are only found at <1% in cyanobacterial genome assemblies. The detection of Proteobacteria can be explained by the presence of these bacteria in cyanobacterial polysaccharide sheaths, which are a source of nutrition for them [24]. Such sheath colonization often hinders efforts to make cyanobacterial cultures axenic [13]. Hence, the complete genome of *Blastomonas* sp., a heterotrophic proteobacterium living in close association with *Cyanobium* sp., has been recently obtained from a non-axenic culture of the latter [25]. Regarding Bacteroidetes, members of this phylum can degrade polymeric compounds, and one isolate was hypothesized to feed on cyanobacterial biomass [46].

**Figure 4:**
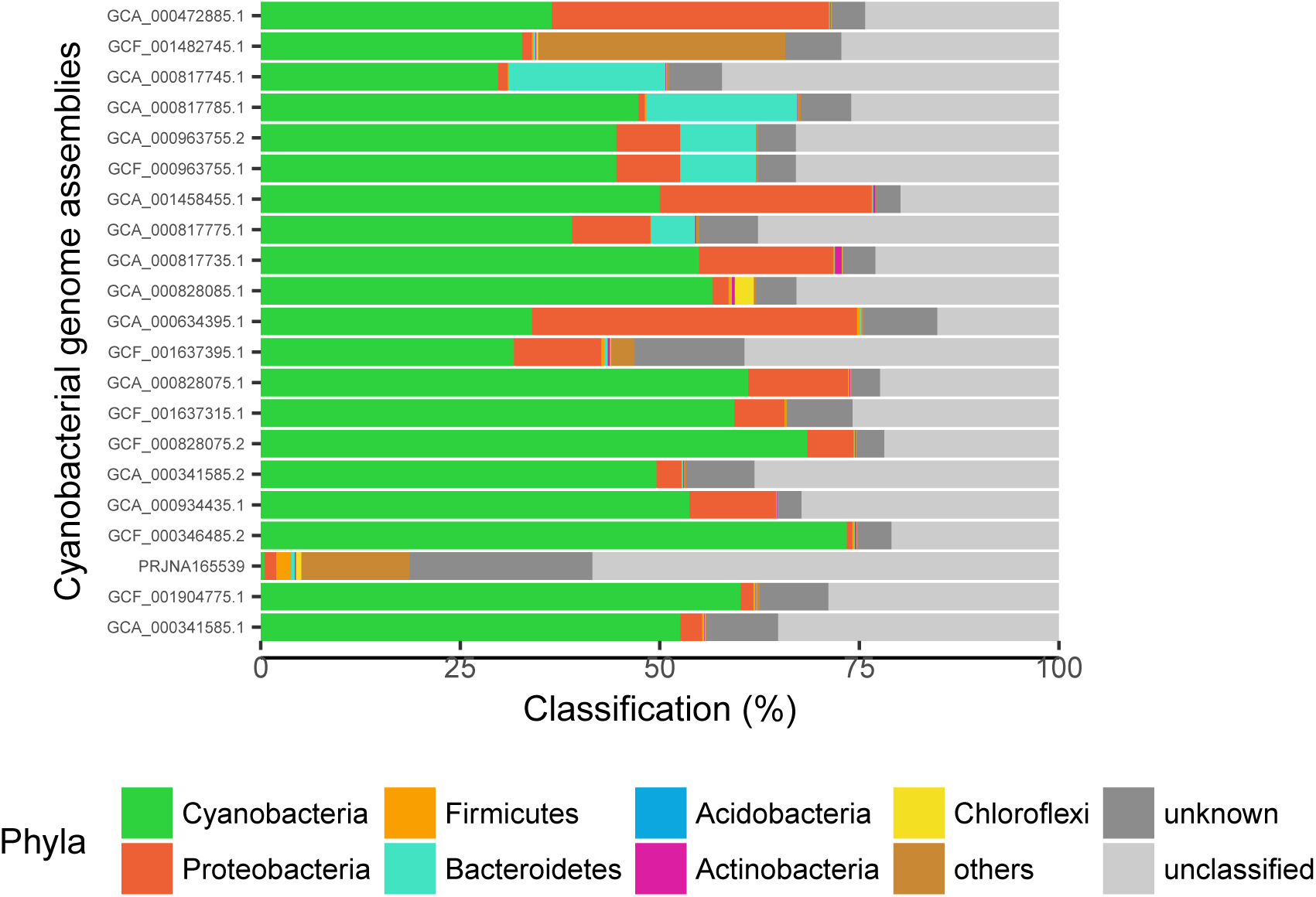
Taxonomic distribution of contaminating sequences in contaminated genomes, based on DIAMOND blastx estimates. The 21 top-ranking contaminated cyanobacterial genome assemblies (see **Table 1**) were analyzed with DIAMOND blastx, as explained in the main text (using a LCA approach against a protein version of our curated Ensembl 30 database). Taxonomic classifications (expressed in *%* of the genomic sequence) are summarized at the phylum level. The “unclassified” classification corresponds to sequences that do not match any reference protein in the database, whereas “unknown” corresponds to high-ranking LCAs (Bacteria or Terrabacteria). “others” include the following phyla (in descending order of frequency): Spirochaetes, Deinococcus-Thermus, Candidatus Tectomicrobia, Verrucomicrobia, Nitrospirae, Tenericutes, Armatimonadetes, Fusobacteria, Thermotogae, Gemmatimonadetes, Synergistetes, Chlamydiae, Thaumarchaeota, Thermodesulfobacteria, Crenarchaeota, Chrysiogenetes, Dictyoglomi. Complete results for the 440 assemblies are available in **Figure S2**.

**Figure 4** further shows that some assemblies have a high level of unclassified sequences (up to 58%). As aforementioned, with methods based on reference databases, the efficiency of classification heavily depends of the diversity of the database. It means that sequences from uncommon organisms, such as extremophiles, are difficult to classify because they are under-represented in reference databases. As an example, *Cyanobacteria bacterium* JGI 0000014-E08 (PRJNA165539; unpublished), which was collected in a lagoon with a reportedly unique microbial diversity (https://www.ncbi.nlm.nih.gov/bioproject/?term=PRJNA165539). displays 58.4% of unclassified sequences. Moreover, a part of the unclassified fraction of this genome is probably of cyanobacterial origin. since cyanobacterial ribosomal gene hits are located on scaffolds with a high level of unclassified segments (**Figure S4**), which also highlights the usefulness of combining the results of multiple methods, especially those of genome-wide approaches that output detailed taxonomic reports (i.e., per scaffold).

### Validation using the sequencing coverage

On average, contaminated genome assemblies have a higher number of scaffolds than non-contaminated assemblies, owing to both the decrease in sequencing coverage and the increase in assembly complexity associated with the mixing of several organisms in the same “genome” (Pearson r = 0.36, P-value = 2.71e-15). In some cases, however, chimerical scaffolds containing a mixture of cyanobacterial and non-cyanobacterial segments can be observed (e.g., GCF_001482745.1, GCA_000643395.1). Confirmed by detailed taxonomic analyses of the scaffolds (**Table S4**), this result suggests that these scaffolds are probably the products of a too greedy concatenation of contigs during the finishing steps of genome sequencing. Ideally, such a bold interpretation should be supported by additional evidence, so as to exclude false positives due to methodological artifacts (e.g., reference database issues, lack of sensitivity of BLAST heuristics) and/or HGT.

To distinguish between genuine contamination and other events, we tried to take advantage of the sequencing coverage, i.e., by monitoring the number of reads mapping at every position of an assembly. Indeed, contaminant DNA is expected to be less abundant than genuine DNA (or at least not in equal proportion), which should result into non-cyanobacterial scaffolds and/or sub-scaffolds having a lower (or at least different) coverage than their cyanobacterial counterparts [31,47]. In principle, the coverage along a genome can easily be obtained by remapping the raw sequencing reads to the corresponding assembly. Unfortunately, a careful search in NCBI SRA for raw sequencing reads corresponding to the 440 cyanobacterial genome assemblies revealed that such data files were only available for 83 (19%) assemblies, of which 63 (14%) were really usable for computing the sequencing coverage. In particular, coverage could be computed for only one (yet the most) highly contaminated assembly (GCA_000472885.1 at rank 1). This large “genome” (15.8 Mbp) is composed of two populations of scaffolds, each one characterized by a distinct value of GC-content (**Figure 5a**). As for most other assemblies of the top-21 (see **Figure S4**), this parameter agrees with our classification of scaffolds as either cyanobacterial (in green) or contaminant (in red), whether using ribosomal genes or DIAMOND blastx for classification. Hence, scaffolds characterized by a high GC-content are those that likely originate from the contaminating organism, rather than from the supposedly sequenced cyanobacterium. As expected, differences in sequencing coverage correlate well with the split into two populations, cyanobacterial scaffolds having a median coverage of 68 (IQR = 9), whereas contaminant scaffolds drop to 26 (IQR = 13). In contrast, when running similar analyses on a non-contaminated cyanobacterial genome assembly (GCF_000214075.1, rank = 379), both the GC-content and the sequencing coverage remain largely uniform, except for plasmid segments visible at extremities of the GC-content range (**Figure 5b**). This confirms that this assembly only contains cyanobacterial scaffolds, as predicted without using the sequencing coverage.

**Figure 5:**
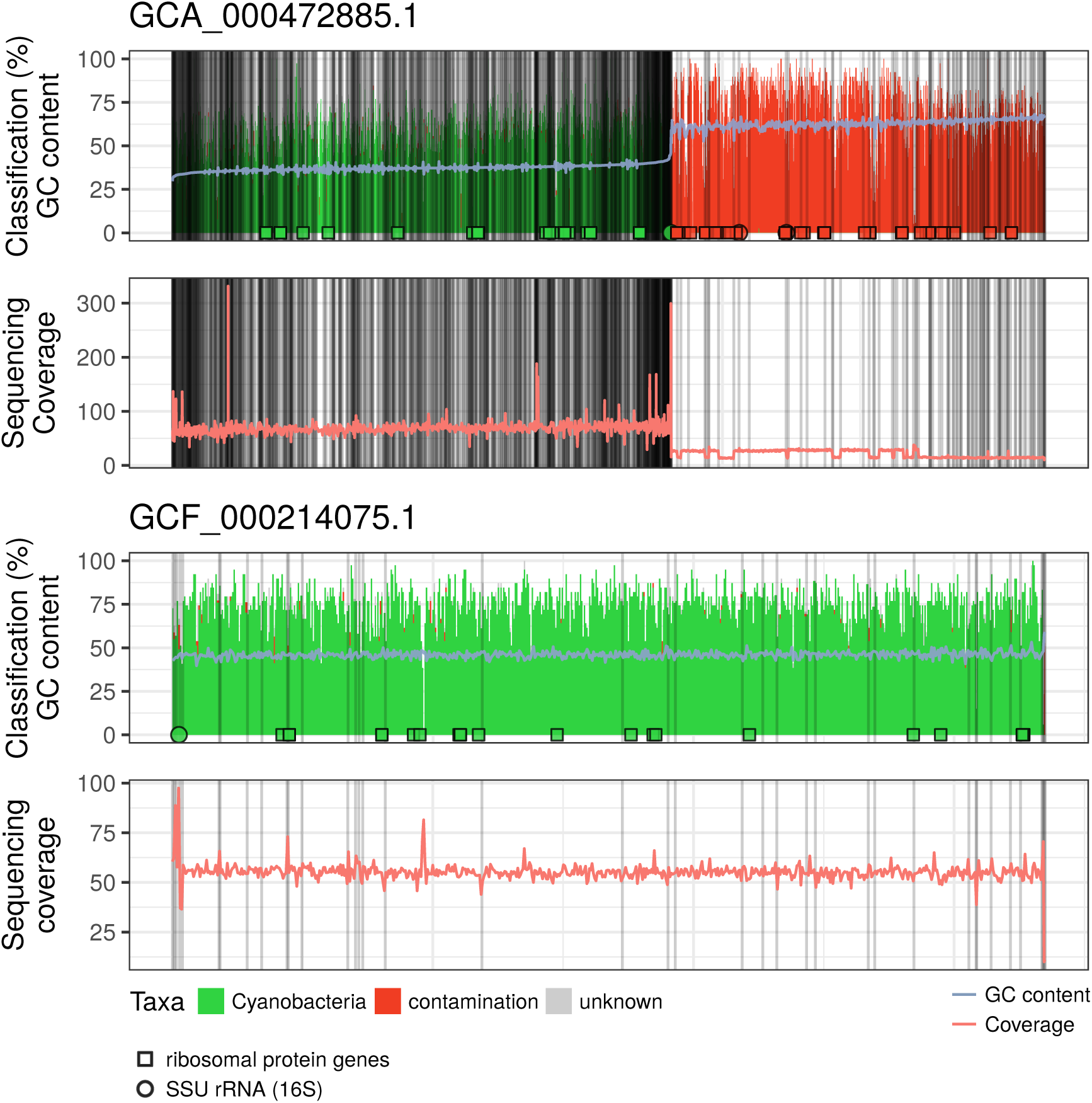
Validation of our methods for detecting contaminants using the sequencing coverage. Two representative cyanobacterial genome assemblies, the highly contaminated GCA_000472885.1 (A) and the non-contaminated GCF_000214075.1 (B), were split into segments of 10,000 nt for detailed analysis. On the X axis, segment-containing scaffolds were sorted by increasing average GC-content, with vertical thin lines representing scaffold boundaries. Top panels: segments were classified as either cyanobacterial (green), contaminated (red), unknown (grey) or unclassified (white), using DIAMOND blastx. The overlaid light blue curve is the GC-content of the corresponding segments. Open symbols give the locations of the detected ribosomal genes (circles: SSU rRNA (16S), squares: ribosomal proteins), either from cyanobacterial sources (green) or from non-cyanobacterial sources (red). Bottom panels: the light red curve gives the sequencing coverage of the segments

For non-cyanobacterial segments in assemblies for which coverage cannot be obtained, it might be difficult to decide between a contamination (i.e., the experimentally-dependent introduction of a foreign DNA sequence in a genome assembly), a HGT event (i.e., the natural introduction of a foreign DNA fragment into a genome to be sequenced) or a methodological artefact (e.g., a chimerical or contaminated genome in the reference database). With respect to artefacts, we are quite confident that they should rarely affect assemblies in our top-21, because of the high level of congruence observed 10 between the detection methods, including published approaches (e.g., CheckM) and in-house protocols based on robust algorithms (e.g., MEGAN-like LCA inference), using a large and curated reference database. Regarding HGT, it is assumed to be common in prokaryotes [48]. However, recent studies do not necessarily agree on the extent of the phenomenon (e.g., [49,50]), and most authors argue that HGT is more frequent among closely related organisms [51–54]. Hence, HGT has been shown to occur within Cyanobacteria [55–58], often at different rates [59,60], depending on the evolutionary constraints acting on the transferred genes, i.e., “stand-alone” functions vs operon-encoded functions [57,61–65]. In the context of the present study, aimed at assessing the contamination level of publicly available genomes, our opinion is that HGT should not be a major issue because we only consider as contaminants the segments that are clearly non-cyanobacterial (i.e., we do not address intra-cyanobacterial contamination) and because real HGT events are more likely to be found in the classified-yet-unknown fraction of the assemblies. Indeed, as HGT makes look highly similar sequences from organisms potentially very distant from a taxonomic point of view, LCA-based methods (Kraken, DIAMOND blastx) label them with high-ranking taxa, such as Bacteria, that our post-processors then convert to “unknown”. Nevertheless, studies specifically designed to assess the extent of HGT in existing genomes should probably double-check that some of their results are not the product of contamination rather than transfer, whether in reference genomes or in genome assemblies under investigation.

## Conclusion

Among the 440 cyanobacterial genome assemblies surveyed in this study, we concluded that 21 were highly contaminated (corresponding to 19 different genomes). Proteobacteria and Bacteroidetes are the most common contaminants, probably due to their close proximity with some Cyanobacteria in the environment, and maybe because of trophic interactions complicating the generation of axenic strains [24]. The six genomes that initially raised our attention, and motivated us to design a consensus pipeline for detecting contaminants, all occupy the top-tier of our global ranking (GCA_000472885.1, rank 1; GCA_000817745.1, rank 3; GCA_000817785.1, rank 6 GCA_000817775.1, rank 8; GCA_000817735.1, rank 9; GCA_000828075.1, rank 13). Moreover, 10 of the top-21 contaminated assemblies discovered in this study have now been removed from RefSeq, or have at least been flagged as suspicious in their metadata by NCBI curators, suggesting that our ranking procedure is accurate.

The vast majority of publicly available cyanobacterial genomes are only slightly contaminated (209 assemblies) or likely non-contaminated (210 assemblies). However, the impact of an error (in particular, a contamination) can be very detrimental [3]. For instance, researchers may infer that a particular enzyme is present in some cyanobacterium, whereas in reality it is absent from the phylum, or that some Cyanobacteria are much more prone to HGT events than others. As researchers, we are more interested by exceptions than by those that play along the rules. Hence, finding an orthologous gene in >400 cyanobacterial genomes attracts less attention (and impact) than a gene specific to Bacteroidetes in a handful of Cyanobacteria. In other words, we are much more prone to ingenuously study erroneous than correct data, thereby amplifying small errors and polluting the scientific literature with incorrect statements resulting from the analysis of contaminated genomes.

It is therefore of prime importance to eliminate as many errors as possible from genomic sequences that now constitute a mainstay of numerous research domains, and this is especially important for newly generated genomes making their way up to reference databases used for taxonomic classification. To this end, releasing the raw reads at the same time as the assembled genome should become mandatory. The availability of such data would allow others not only to repeat (or improve) the assembly process, but also spot obvious contaminants based on discrepancies in sequencing coverage. However, such a policy would likely be insufficient, because one cannot expect every genomicist to look for contaminants in tens of thousand of bacterial genomes by downloading and re-mapping raw reads on the corresponding assemblies. To ease the task of the large scientific community using genomic data, the sequencing coverage of each position, as well as other key statistics obtained during the assembly process (e.g., the number of non-mapping reads, the number of incongruently mapping paired ends) should be published along with the genome assembly. This would allow researchers to discard from their homology searches all the regions with a low coverage (probable contaminants) or with a high coverage (probable repeated elements). Technically, making available the .bam files in addition to the raw read files would provide most of the information needed by others. Implementing this constraint is then simply a matter of editorial policy, exactly as when, decades ago, *Molecular Biology and Evolution* decided to reject any manuscript featuring a phylogenetic tree devoid of statistical support (e.g., bootstrap). In contrast, factoring genomic meta-data (sequencing coverage, paired-end incongruence, etc.) into homology searches is likely to be more difficult. Indeed, it will require devising a standard format and modifying numerous software packages to take advantage of it, but it is certainly worth the effort if we want to make the most of genomic data.

## Materials and Methods

### Cyanobacterial genome assemblies

Cyanobacterial genome assemblies have been collected from four databases using a custom Perl script. The latter uses the assembly number to query public sequence databases according to a predetermined order, so that the genome redundancy across banks is handled consistently. Genomes were first collected from the release 30 of Ensembl Bacteria [42], then from the NCBI (with a priority on RefSeq [66], then from PATRIC [67], and finally from the bacterial part of the JGI genome portal [68]). A first batch of downloads was carried out on the 14th of April 2016 and a second batch on the 11th of December 2016.

Assemblies were first analyzed using QUAST 2.3 [2] with default values. The number of scaffolds, the number of scaffolds >1000 nt, the total length of the genomes, the total length of the genomes based on scaffolds >1000 nt, the length of the largest scaffold, the GC-content, the N50/N75 values, the L50/L75 values and the number of N’s per 100 kbp were collected and summarized in **Table S1**, along with the morphology, the habitat and the taxonomy of each corresponding strain.

### SSU rRNA (16S) analyses

SSU rRNA (16S) genes in cyanobacterial genome assemblies were predicted with RNAmmer 1.2, a rRNA predictor using hidden Markov models [28]. The putative extracted SSU rRNA sequences were taxonomically classified by SINA 1.2.11, a multiple sequence alignment and classifier tool (associated with SILVA rRNA database) [29]. SINA was used in combination with the non-redundant and curated release 123.1 of SILVA, which contains 645,151 SSU rRNA (16S) sequences for the classification [36]. A rough estimate of the contamination level in each assembly was computed by taking the ratio of the number of SSU rRNA (16S) genes classified to foreign (i.e., non-cyanobacterial) Bacteria to the total number of predicted SSU rRNA (16S) genes.

The distance matrix used for benchmarking reference databases was computed by aligning the SSU rRNA (16S) sequences with MAFFT 7.273 [69], then selecting the conserved positions of the alignment with BMGE 1.12 [70] and finally calculating the distance matrix with IQPNNI 3.3.2, a tree-reconstruction algorithm based on maximum likelihood (ML) and used here with the GTR+Γ4 model [71].

### Ribosomal protein analyses

The contamination level estimates of cyanobacterial genome assemblies were refined using a broadly-sampled dataset of 52 multiple sequence alignments (MSAs) of ribosomal proteins aligned with MAFFT. The dataset was assembled from a ribosomal protein database for prokaryotes, RiboDB 1.4.0 [39], available at https://ribodb.univ-lyon1.fr/. These protein sequences were used as a reference to detect and classify contaminating sequences with “42”, a program whose aim is to add (and to optionally align) sequences to a pre-existing MSA while controlling for orthology relationships (Baurain et al., to be published elsewhere; https://bitbucket.org/dbaurain/42/). Here, we used “42”′s ability to spot contaminants by attempting to add ribosomal proteins from each of our 440 cyanobacterial genomes to each MSA of our dataset of ribosomal MSAs. In practice, “42” searches each genome for sequences that are homologous to sequences of the current MSA and then, through advanced heuristics, sort out orthologues from possible paralogues. Each orthologous sequence is further classified by computing the LCA of the three sequences of the MSA that are the most similar to the newly added orthologue, with a minimum identity threshold of 70% and a minimum alignment length threshold of 30 amino acids. By default, we expect that the inferred LCA is in agreement with the taxonomy of the genome from which comes the orthologue, i.e., of cyanobacterial origin. If not, the orthologue is considered as a contaminant of the genome assembly (or remains unclassified if not matching anything close enough in the reference database).

We used RAxML trees inferred under the LG+F model [72] and the CAT approximation to check whether reference Cyanobacteria were monophyletic for all 52 ribosomal proteins. When automated parsing of the bipartitions suggested that it was not the case, visual inspection of the ML tree allowed us to decide between HGT (6 genes) and sequence divergence/reconstruction error (3 genes) as the cause for the non-monophyly. Before proceeding, a few xenologous sequences were thus removed from 6 MSAs (L12, L31, L36, S16, S21, uS4). Moreover, 6 MSAs (L10, L25, L28, L29, L32, S21) were completely discarded due to poor performance of “42” (i.e., no or too few added sequences, orthology assessment issues, for further details, see **Table S5**). The remaining 46 MSAs represented 178,634 sequences sampled across a variety of 3474 different bacterial and archaeal strains in total. They were used to estimate the contamination levels as with SSU rRNA (16S) genes, but by summing over all individual ribosomal proteins found in each cyanobacterial genome assembly, i.e., ignoring their possible collocation on the chromosome.

### CheckM analyses

We analysed the 440 cyanobacterial genome assemblies with CheckM 1.0.7. CheckM uses a concatenation of predicted ribosomal proteins to automatically place the assembly under study in a reference genome tree. Then, it estimates the completeness and the contamination level of the assembly by searching the sequence for lineage-specific marker genes provided with the software (see [14]). We used the typical automatical workflow option *lineage*_*wf.* The contamination percentage extracted from the output was directly considered as the contamination level estimate.

### Kraken analyses

Kraken 0.10.5-beta is a program that assigns taxonomic labels to genomic DNA sequences [30]. It uses exact (signature) kmer (21, 25, 31 nt) alignment to associate a taxonomic tree with a kmer database that it builds itself according to a default or user-specified genome set [24]. We used a newly created curated database derived from Ensembl Bacteria 30 (see **Supplemental Note 1** for details) with three different confidence thresholds (0.02, 0.04 and 0.06) to analyze cyanobacterial pseudo-reads. Kraken labels were further post-processed to classify pseudo-reads as either cyanobacterial, contaminating (non-cyanobacterial), unknown (too high-ranking LCA, such as “Bacteria” or “Terrabacteria”) or unclassified (no hit in the database). Finally, a global contamination level was computed for each assembly by taking the ratio of the number of non-cyanobacterial pseudo-reads to the total number of pseudo-reads.

### DIAMOND blastx analyses

To check the effect of using proteins instead of whole nucleotide genomes on detection sensitivity, we repeated most Kraken analyses with DIAMOND 0.8.22.84 blastx [32]. For this, we built a DIAMOND database from the corresponding 27,762 complete proteomes of our curated Ensembl 30 genome set. Cyanobacterial genome assemblies, still split into pseudo-reads of 250 nt, were BLASTed against this protein database. Each pseudo-read was then labelled by computing the LCA of its best hits (excluding self-matches), provided they had a bit-score ≥80 and within 95% of the bit-score of the first hit (MEGAN-like algorithm [44]). We tested three different thresholds for the minimal number of hits required for LCA inference (1, 2 or 3) and chose to use a minimal number of 1 hit. From there, pseudo-read classification and contamination level estimation were carried out exactly as for Kraken. Decontaminated cyanobacterial genome assemblies, in which regions corresponding to contaminating pseudo-reads have been removed (or only masked with N’s), are available for download at https://figshare.com/s/08193cb71f0a65d99c69.

### CONCOCT analyses

We also analyzed cyanobacterial genome assemblies with CONCOCT 0.4.1 [31] using three default values for kmer size (4, 5 and 6). For each assembly, the script cut_up_fasta.py (provided with CONCOCT) was used to cut the genome sequences into non-overlapping segments of 10,000 nt. Dummy coverage files with a value of 1 for every segment were also created. CONCOCT was used on the resulting files with default values: a length threshold of 1000 nt for the segments and a number of groups of 400 for the clustering based on the principal component analysis (PCA) of kmer frequencies. For each assembly, we computed the total number of segments for the whole assembly and counted the number of segments associated with every group determined by CONCOCT. Then, we selected the group containing the largest number of segments and considered it to be the “core” genome without any contamination or recent HGT. To estimate the contamination level of an assembly, we used the simple following formula: one minus the ratio between the number of segments belonging to the largest group and the total number of segments.

Importantly, CONCOCT was designed for metagenomic analyses and requires coverage data to realize a meaningful clustering of sequencing reads after the PCA [31]. This is also the case for most other programs created for metagenomics, such as MetaBat [47], VizBin [73], GroopM [74], MaxBin [75] and the older CompostBin [76]. Unfortunately, sequencing coverage could be obtained for only 63 assemblies (see *Validation using the sequencing coverage* for details). Yet, when we reran CONCOCT using coverage files, we did not observe any large difference in clustering with respect to the results obtained without coverage (data not shown). However, none of these 63 assemblies (including 7 from the 97 atypical assemblies), except*Mastigocoleus testarum* BC008 (GCA_000472885.1, unpublished), were highly contaminated (DIAMOND blastx level <2%). Thus, the use of real coverage values might have had little effect on the clustering results. This interpretation is strengthened by the observation that real coverage values were indeed relatively constant within each of the 62 non-contaminated assemblies, as assessed through the corresponding quartile coefficients of dispersion (median = 0.055, IQR = 0.053).

### Global ranking

Since all methods were in general agreement, we decided to retain all of them in our final global ranking of cyanobacterial genome assemblies. This consensus ranking was computed by averaging the ranks of each assembly in the individual rankings obtained for each of the six methods (**Table S2**). For the 170 assemblies included in the reference genome database, Kraken results were considered as missing values (NA) when taking the rank average. Spearman rank correlations were computed for all methods and the global ranking, considering all 440 assemblies, only the 343 typical assemblies, or only the 50 top-ranking assemblies out of the 440 (**Table S3**), using the R option pairwise.complete.obs to handle missing values.

### Validation using the sequencing coverage

To confirm that foreign sequences in contaminated cyanobacterial genomes were genuine contaminants, we checked that their sequencing coverage was different from the coverage of cyanobacterial sequences. Coverage was estimated using BBMap (http://bbmap.sourceforge.net/) from raw read data (when available). However, this was possible for only 63 of the 83 available SRA files: 12 genomes had *avf*_*fold* values <5 and 8 genomes returned no result at all. Among the 63 assemblies with coverage, only one (GCA_000472885.1) was part of our top-21 ranking, all other assemblies corresponding to slightly or non-contaminated genomes.

For each assembly, sequencing coverage, GC-content (computed with a custom Perl script) and taxonomic classification (cyanobacterial, contaminated, unknown and unclassified sequence fractions, as determined by DIAMOND blastx) were computed on 10,000-nt segments (as for CONCOCT) and used for generating combined genome maps. In the corresponding plots, scaffolds are arranged by GC-content, which allows highlighting the covariation of the latter with contamination level and sequencing coverage. As an additional congruence test, the positions of the SSU rRNA (16S) and ribosomal protein genes classified to Cyanobacteria and/or to contaminant organisms were also plotted below their corresponding scaffolds.

## Author Statements

## Acknowledgment

We thank Yoan Bouzin for his help in elaborating the downloading script and Rosa Gago for her artistic contribution in the graphical abstract.

## Author contributions

LC and DB designed the analyses, LM performed the SSU rRNA (16S) analyses and drew most of the figures, MVV performed the ribosomal protein analyses and designed the graphical abstract, RRL performed the CONCOCT analyses, LC performed all other analyses and drew the remaining figures, BD, YL and LC collected the details on the cyanobacterial strains, AM, BD and YL prototyped some aspects of the Kraken analyses, DS helped with the computer cluster and the statistics, HP suggested additional analyses and helped structuring the manuscript, LC and DB wrote the manuscript, with the assistance of AW, LM, MVV, YL, HP and EJJ. All authors read and approved the final manuscript.

## Ethics approval and consent to participate

Not applicable.

## Consent for publication

Not applicable.

## Data summary

The datasets generated and/or analyzed during the current study are available at:

- https://figshare.com/s/bdcc314a7b90b00c1274
- https://figshare.com/s/08193cb71f0a65d99c69
- https://figshare.com/s/344be979f5f2237ad735

## References

1. Simão FA, Waterhouse RM, Ioannidis P, Kriventseva EV, Zdobnov EM. BUSCO: assessing genome assembly and annotation completeness with single-copy orthologs. Bioinformatics. 2015;31: 3210–3212. doi:10.1093/bioinformatics/btv351

2. Gurevich A, Saveliev V, Vyahhi N, Tesler G. QUAST: quality assessment tool for genome assemblies. Bioinformatics. 2013;29: 1072–1075. doi:10.1093/bioinformatics/btt086

3. Kozlov AM, Zhang J, Yilmaz P, Glöckner FO, Stamatakis A. Phylogeny-aware identification and correction of taxonomically mislabeled sequences. Nucleic Acids Res. 2016; gkw396. doi:10.1093/nar/gkw396

4. Ballenghien M, Faivre N, Galtier N. Patterns of cross-contamination in a multispecies population genomic project: detection, quantification, impact, and solutions. BMC Biol. 2017;15: 25. doi:10.1186/s12915-017-0366-6

5. Merchant S, Wood DE, Salzberg SL. Unexpected cross-species contamination in genome sequencing projects. PeerJ. 2014;2: e675. doi:10.7717/peerj.675

6. Simion P, Philippe H, Baurain D, Jager M, Richter DJ, Franco AD, et al. A Large and Consistent Phylogenomic Dataset Supports Sponges as the Sister Group to All Other Animals. Curr Biol. 2017;27: 958–967. doi:10.1016/j.cub.2017.02.031

7. Finet C, Timme RE, Delwiche CF, Marlétaz F. Multigene Phylogeny of the Green Lineage Reveals the Origin and Diversification of Land Plants. Curr Biol. 2010;20: 2217–2222. doi:10.1016/j.cub.2010.11.035

8. Laurin-Lemay S, Brinkmann H, Philippe H. Origin of land plants revisited in the light of sequence contamination and missing data. Curr Biol. 2012;22: R593–R594. doi:10.1016/j.cub.2012.06.013

9. Schierwater B, Kolokotronis S-O, Eitel M, DeSalle R. The Diploblast-Bilateria sister hypothesis. Commun Integr Biol. 2009;2: 403–405. doi:10.4161/cib.2.5.8763

10. Philippe H, Brinkmann H, Lavrov DV, Littlewood DTJ, Manuel M, Wörheide G, et al. Resolving Difficult Phylogenetic Questions: Why More Sequences Are Not Enough. PLOS Biol. 2011;9: e1000602. doi:10.1371/journal.pbio.1000602

11. Longo MS, O’Neill MJ, O’Neill RJ. Abundant Human DNA Contamination Identified in Non-Primate Genome Databases. PLOS ONE. 2011;6: e16410. doi:10.1371/journal.pone.0016410

12. Philippe H, Vienne DM de, Ranwez V, Roure B, Baurain D, Delsuc F. Pitfalls in supermatrix phylogenomics. Eur J Taxon. 2017;0. doi:10.5852/ejt.2017.283

13. Rippka R, Deruelles J, Waterbury JB, Herdman M, Stanier RY. Generic Assignments, Strain Histories and Properties of Pure Cultures of Cyanobacteria. Microbiology. 1979;111: 1–61. doi:10.1099/00221287-111-1-1

14. Parks DH, Imelfort M, Skennerton CT, Hugenholtz P, Tyson GW. CheckM: assessing the quality of microbial genomes recovered from isolates, single cells, and metagenomes. Genome Res. 2015;25: 1043–1055. doi:10.1101/gr.186072.114

15. Tennessen K, Andersen E, Clingenpeel S, Rinke C, Lundberg DS, Han J, et al. ProDeGe: a computational protocol for fully automated decontamination of genomes. ISME J. 2016;10: 269–272. doi:10.1038/ismej.2015.100

16. Lux M, Krüger J, Rinke C, Maus I, Schlüter A, Woyke T, et al. acdc – Automated Contamination Detection and Confidence estimation for single-cell genome data. BMC Bioinformatics. 2016;17: 543. doi:10.1186/s12859-016-1397-7

17. Alvarenga DO, Fiore MF, Varani AM. A Metagenomic Approach to Cyanobacterial Genomics. Front Microbiol. 2017;8. doi:10.3389/fmicb.2017.00809

18. Skoglund P, Northoff BH, Shunkov MV, Derevianko AP, Pääbo S, Krause J, et al. Separating endogenous ancient DNA from modern day contamination in a Siberian Neandertal. Proc Natl Acad Sci. 2014;111: 2229–2234. doi:10.1073/pnas.1318934111

19. Brown MW, Heiss AA, Kamikawa R, Inagaki Y, Yabuki A, Tice AK, et al. Phylogenomics Places Orphan Protistan Lineages in a Novel Eukaryotic Super-Group. Genome Biol Evol. 2018;10: 427–433. doi:10.1093/gbe/evy014

20. Knoll AH. The geological consequences of evolution. Geobiology. 2003;1: 3–14. doi:10.1046/j.1472-4669.2003.00002.x

21. Kopp RE, Kirschvink JL, Hilburn IA, Nash CZ. The Paleoproterozoic snowball Earth: A climate disaster triggered by the evolution of oxygenic photosynthesis. Proc Natl Acad Sci U S A. 2005;102: 11131–11136. doi:10.1073/pnas.0504878102

22. Ochoa de Alda JAG, Esteban R, Diago ML, Houmard J. The plastid ancestor originated among one of the major cyanobacterial lineages. Nat Commun. 2014;5: 4937. doi:10.1038/ncomms5937

23. Whitton BA, Potts M. Introduction to the Cyanobacteria. In: Whitton BA, editor. Ecology of Cyanobacteria II. Springer Netherlands; 2012. pp. 1–13. Available: http://link.springer.com/chapter/10.1007/978-94-007-3855-3_1

24. Stuart RK, Mayali X, Lee JZ, Craig Everroad R, Hwang M, Bebout BM, et al. Cyanobacterial reuse of extracellular organic carbon in microbial mats. ISME J. 2016;10: 1240–1251. doi:10.1038/ismej.2015.180

25. Lima ARJ, Siqueira AS, Santos BGS dos, Silva FDF da, Lima CP, Cardoso JF, et al. Draft Genome Sequence of Blastomonas sp. Strain CACIA 14H2, a Heterotrophic Bacterium Associated with Cyanobacteria. Genome Announc. 2014;2: e01200–13. doi:10.1128/genomeA.01200-13

26. Lee JZ, Burow LC, Woebken D, Everroad RC, Kubo MD, Spormann AM, et al. Fermentation couples Chloroflexi and sulfate-reducing bacteria to Cyanobacteria in hypersaline microbial mats. Microb Physiol Metab. 2014;5: 61. doi:10.3389/fmicb.2014.00061

27. Peeters K, Verleyen E, Hodgson DA, Convey P, Ertz D, Vyverman W, et al. Heterotrophic bacterial diversity in aquatic microbial mat communities from Antarctica. Polar Biol. 2012;35: 543–554. doi:10.1007/s00300-011-1100-4

28. Lagesen K, Hallin P, Rødland EA, Stærfeldt H-H, Rognes T, Ussery DW. RNAmmer: consistent and rapid annotation of ribosomal RNA genes. Nucleic Acids Res. 2007;35: 3100–3108. doi:10.1093/nar/gkm160

29. Pruesse E, Peplies J, Glöckner FO. SINA: Accurate high-throughput multiple sequence alignment of ribosomal RNA genes. Bioinformatics. 2012;28: 1823–1829. doi:10.1093/bioinformatics/bts252

30. Wood DE, Salzberg SL. Kraken: ultrafast metagenomic sequence classification using exact alignments. Genome Biol. 2014;15: R46. doi:10.1186/gb-2014-15-3-r46

31. Alneberg J, Bjarnason BS, de Bruijn I, Schirmer M, Quick J, Ijaz UZ, et al. Binning metagenomic contigs by coverage and composition. Nat Methods. 2014;11: 1144–1146. doi:10.1038/nmeth.3103

32. Buchfink B, Xie C, Huson DH. Fast and sensitive protein alignment using DIAMOND. Nat Methods. 2015;12: 59–60. doi:10.1038/nmeth.3176

33. Komárek J. A polyphasic approach for the taxonomy of cyanobacteria: principles and applications. Eur J Phycol. 2016;51: 346–353. doi:10.1080/09670262.2016.1163738

34. Ponce-Toledo RI, Deschamps P, López-García P, Zivanovic Y, Benzerara K, Moreira D. An Early-Branching Freshwater Cyanobacterium at the Origin of Plastids. Curr Biol. 2017;27: 386–391. doi:10.1016/j.cub.2016.11.056

35. Woese CR, Fox GE, Zablen L, Uchida T, Bonen L, Pechman K, et al. Conservation of primary structure in 16S ribosomal RNA. Nature. 1975;254: 83–86. doi:10.1038/254083a0

36. Quast C, Pruesse E, Yilmaz P, Gerken J, Schweer T, Yarza P, et al. The SILVA ribosomal RNA gene database project: improved data processing and web-based tools. Nucleic Acids Res. 2013;41: D590–D596. doi:10.1093/nar/gks1219

37. Klappenbach JA, Dunbar JM, Schmidt TM. rRNA Operon Copy Number Reflects Ecological Strategies of Bacteria. Appl Environ Microbiol. 2000;66: 1328–1333. doi:10.1128/AEM.66.4.1328-1333.2000

38. Engene N, Gerwick WH. Intra-genomic 16S rRNA gene heterogeneity in cyanobacterial genomes. Fottea. 2011;11: 17–24. doi:10.5507/fot.2011.003

39. Jauffrit F, Penel S, Delmotte S, Rey C, Vienne DM de, Gouy M, et al. RiboDB Database: A Comprehensive Resource for Prokaryotic Systematics. Mol Biol Evol. 2016; msw088. doi:10.1093/molbev/msw088

40. Khayrullina GA, Raabe CA, Hoe CH, Becker K, Reinhardt R, Tang TH, et al. Transcription Analysis and Small Non-Protein Coding RNAs Associated with Bacterial Ribosomal Protein Operons. Curr Med Chem. 2012;19: 5187–5198. doi:10.2174/092986712803530485

41. Brochier C, Philippe H, Moreira D. The evolutionary history of ribosomal protein RpS14: Trends Genet. 2000;16: 529–533. doi:10.1016/S0168-9525(00)02142-9

42. Aken BL, Ayling S, Barrell D, Clarke L, Curwen V, Fairley S, et al. The Ensembl gene annotation system. Database. 2016;2016: baw093. doi:10.1093/database/baw093

43. Boratyn GM, Schäffer AA, Agarwala R, Altschul SF, Lipman DJ, Madden TL. Domain enhanced lookup time accelerated BLAST. Biol Direct. 2012;7: 12. doi:10.1186/1745-6150-7-12

44. Huson DH, Auch AF, Qi J, Schuster SC. MEGAN analysis of metagenomic data. Genome Res. 2007;17: 377–386. doi:10.1101/gr.5969107

45. Darriba D, Flouri T, Stamatakis A. The state of software for evolutionary biology. Mol Biol Evol. doi:10.1093/molbev/msy014

46. Ben Hania W, Joseph M, Bunk B, Spröer C, Klenk H-P, Fardeau M-L, et al. Characterization of the first cultured representative of a Bacteroidetes clade specialized on the scavenging of cyanobacteria. Environ Microbiol. 2017;19: 1134–1148. doi:10.1111/1462-2920.13639

47. Kang DD, Froula J, Egan R, Wang Z. MetaBAT, an efficient tool for accurately reconstructing single genomes from complex microbial communities. PeerJ. 2015;3: e1165. doi:10.7717/peerj.1165

48. Doolittle WF. Phylogenetic Classification and the Universal Tree. Science. 1999;284: 2124–2128. doi:10.1126/science.284.5423.2124

49. Andam CP, Fournier GP, Gogarten JP. Multilevel populations and the evolution of antibiotic resistance through horizontal gene transfer. FEMS Microbiol Rev. 2011;35: 756–767. doi:10.1111/j.1574-6976.2011.00274.x

50. Wiedenbeck J, Cohan FM. Origins of bacterial diversity through horizontal genetic transfer and adaptation to new ecological niches. FEMS Microbiol Rev. 2011;35: 957–976. doi:10.1111/j.1574-6976.2011.00292.x

51. Manen J-F, Falquet J. The cpcB-cpcA locus as a tool for the genetic characterization of the genus Arthrospira (Cyanobacteria): evidence for horizontal transfer. Int J Syst Evol Microbiol. 2002;52: 861–867. doi:10.1099/00207713-52-3-861

52. Papke RT, Zhaxybayeva O, Feil EJ, Sommerfeld K, Muise D, Doolittle WF. Searching for species in haloarchaea. Proc Natl Acad Sci. 2007;104: 14092–14097. doi:10.1073/pnas.0706358104

53. Popa O, Hazkani-Covo E, Landan G, Martin W, Dagan T. Directed networks reveal genomic barriers and DNA repair bypasses to lateral gene transfer among prokaryotes. Genome Res. 2011;21: 599–609. doi:10.1101/gr.115592.110

54. Popa O, Landan G, Dagan T. Phylogenomic networks reveal limited phylogenetic range of lateral gene transfer by transduction. ISME J. 2017;11: 543–554. doi:10.1038/ismej.2016.116

55. Zhaxybayeva O. Phylogenetic analyses of cyanobacterial genomes: Quantification of horizontal gene transfer events. Genome Res. 2006;16: 1099–1108. doi:10.1101/gr.5322306

56. Shi T, Falkowski PG. Genome evolution in cyanobacteria: The stable core and the variable shell. Proc Natl Acad Sci. 2008;105: 2510–2515. doi:10.1073/pnas.0711165105

57. Tooming-Klunderud A, Sogge H, Rounge TB, Nederbragt AJ, Lagesen K, Glöckner G, et al. From Green to Red: Horizontal Gene Transfer of the Phycoerythrin Gene Cluster between Planktothrix Strains. Appl Environ Microbiol. 2013;79: 6803–6812. doi:10.1128/AEM.01455-13

58. Khan MA, Mahmudi O, Ullah I, Arvestad L, Lagergren J. Probabilistic inference of lateral gene transfer events. BMC Bioinformatics. 2016;17: 431. doi:10.1186/s12859-016-1268-2

59. Rivera MC, Jain R, Moore JE, Lake JA. Genomic evidence for two functionally distinct gene classes. Proc Natl Acad Sci. 1998;95: 6239–6244.

60. Jain R, Rivera MC, Lake JA. Horizontal gene transfer among genomes: The complexity hypothesis. Proc Natl Acad Sci. 1999;96: 3801–3806. doi:10.1073/pnas.96.7.3801

61. Rudi K, Skulberg OM, Jakobsen KS. Evolution of Cyanobacteria by Exchange of Genetic Material among Phyletically Related Strains. J Bacteriol. 1998;180: 3453–3461.

62. Mikalsen B, Boison G, Skulberg OM, Fastner J, Davies W, Gabrielsen TM, et al. Natural Variation in the Microcystin Synthetase Operon mcyABC and Impact on Microcystin Production in Microcystis Strains. J Bacteriol. 2003;185: 2774–2785. doi:10.1128/JB.185.9.2774-2785.2003

63. Klassen JL. Pathway Evolution by Horizontal Transfer and Positive Selection Is Accommodated by Relaxed Negative Selection upon Upstream Pathway Genes in Purple Bacterial Carotenoid Biosynthesis. J Bacteriol. 2009;191: 7500–7508. doi:10.1128/JB.01060-09

64. Martiny AC, Huang Y, Li W. Occurrence of phosphate acquisition genes in Prochlorococcus cells from different ocean regions. Environ Microbiol. 2009;11: 1340–1347. doi:10.1111/j.1462-2920.2009.01860.x

65. Tooming-Klunderud A, Fewer DP, Rohrlack T, Jokela J, Rouhiainen L, Sivonen K, et al. Evidence for positive selection acting on microcystin synthetase adenylation domains in three cyanobacterial genera. BMC Evol Biol. 2008;8: 256. doi:10.1186/1471-2148-8-256

66. O’Leary NA, Wright MW, Brister JR, Ciufo S, Haddad D, McVeigh R, et al. Reference sequence (RefSeq) database at NCBI: current status, taxonomic expansion, and functional annotation. Nucleic Acids Res. 2016;44: D733–745. doi:10.1093/nar/gkv1189

67. Wattam AR, Abraham D, Dalay O, Disz TL, Driscoll T, Gabbard JL, et al. PATRIC, the bacterial bioinformatics database and analysis resource. Nucleic Acids Res. 2013; gkt1099. doi:10.1093/nar/gkt1099

68. Nordberg H, Cantor M, Dusheyko S, Hua S, Poliakov A, Shabalov I, et al. The genome portal of the Department of Energy Joint Genome Institute: 2014 updates. Nucleic Acids Res. 2014;42: D26–31. doi:10.1093/nar/gkt1069

69. Katoh K, Standley DM. MAFFT Multiple Sequence Alignment Software Version 7: Improvements in Performance and Usability. Mol Biol Evol. 2013;30: 772–780. doi:10.1093/molbev/mst010

70. Criscuolo A, Gribaldo S. Large-Scale Phylogenomic Analyses Indicate a Deep Origin of Primary Plastids within Cyanobacteria. Mol Biol Evol. 2011;28: 3019–3032. doi:10.1093/molbev/msr108

71. Vinh LS, Haeseler A von. IQPNNI: Moving Fast Through Tree Space and Stopping in Time. Mol Biol Evol. 2004;21: 1565–1571. doi:10.1093/molbev/msh176

72. Stamatakis A. RAxML version 8: a tool for phylogenetic analysis and post-analysis of large phylogenies. Bioinformatics. 2014;30: 1312–1313. doi:10.1093/bioinformatics/btu033

73. Laczny CC, Sternal T, Plugaru V, Gawron P, Atashpendar A, Margossian HH, et al. VizBin - an application for reference-independent visualization and human-augmented binning of metagenomic data. Microbiome. 2015;3: 1. doi:10.1186/s40168-014-0066-1

74. Imelfort M, Parks D, Woodcroft BJ, Dennis P, Hugenholtz P, Tyson GW. GroopM: an automated tool for the recovery of population genomes from related metagenomes. PeerJ. 2014;2: e603. doi:10.7717/peerj.603

75. Wu Y-W, Tang Y-H, Tringe SG, Simmons BA, Singer SW. MaxBin: an automated binning method to recover individual genomes from metagenomes using an expectation-maximization algorithm. Microbiome. 2014;2: 26. doi:10.1186/2049-2618-2-26

76. Chatterji S, Yamazaki I, Bai Z, Eisen JA. CompostBin: A DNA Composition-Based Algorithm for Binning Environmental Shotgun Reads. In: Vingron M, Wong L, editors. Research in Computational Molecular Biology. Springer Berlin Heidelberg; 2008. pp. 17–28. Available: http://link.springer.com/chapter/10.1007/978-3-540-78839-3_3

